# Characterizing directional dynamics of semantic prediction based on inter-regional temporal generalization

**DOI:** 10.1101/2024.02.13.580183

**Authors:** Fahimeh Mamashli, Sheraz Khan, Elaheh Hatamimajoumerd, Mainak Jas, Işıl Uluç, Kaisu Lankinen, Jonas Obleser, Angela D. Friederici, Burkhard Maess, Jyrki Ahveninen

## Abstract

The event-related potential/field component N400(m) has been widely used as a neural index for semantic prediction. It has long been hypothesized that feedback information from inferior frontal areas plays a critical role in generating the N400. However, due to limitations in causal connectivity estimation, direct testing of this hypothesis has remained difficult. Here, magnetoencephalography (MEG) data was obtained during a classic N400 paradigm where the semantic predictability of a fixed target noun was manipulated in simple German sentences. To estimate causality, we implemented a novel approach based on machine learning and temporal generalization to estimate the effect of inferior frontal gyrus (IFG) on temporal areas. In this method, a support vector machine (SVM) classifier is trained on each time point of the neural activity in IFG to classify less predicted (LP) and highly predicted (HP) nouns and then tested on all time points of superior/middle temporal sub- regions activity (and vice versa, to establish spatio-temporal evidence for or against causality). The decoding accuracy was significantly above chance level when the classifier was trained on IFG activity and tested on future activity in superior and middle temporal gyrus (STG/MTG). The results present new evidence for a model predictive speech comprehension where predictive IFG activity is fed back to shape subsequent activity in STG/MTG, implying a feedback mechanism in N400 generation. In combination with the also observed strong feedforward effect from left STG/MTG to IFG, our findings provide evidence of dynamic feedback and feedforward influences between IFG and temporal areas during N400 generation.

## Introduction

Lexical semantic prediction has been associated with an event-related response component termed N400 (Kutas and Hillyard, 1980, 1989; Federmeier, 2007; Lau et al., 2009; Lau et al., 2013b; Lau et al., 2013a). N400 amplitude is sensitive to the previous context, and its amplitude is reliably reduced following a supportive or predictive context (Kutas and Hillyard, 1984; Federmeier et al., 2007; Kutas and Federmeier, 2011; Wlotko and Federmeier, 2012).

Using MEG, several approaches have been taken to identify the brain network underlying the N400m, the magnetic counterpart of N400 observed in EEG. Generally, the results suggest left hemispheric dominance and involvement of temporal and inferior frontal sources in N400(m) generation (Halgren et al., 2002; Marinkovic et al., 2003; Pulvermüller et al., 2005; Maess et al., 2006; Pylkkanen and McElree, 2007; Salmelin, 2007; Dikker and Pylkkanen, 2012). Despite numerous studies on N400, the information flow between regions appearing to contribute to N400 generation has remained elusive.

Theoretical models on language processing suggest that superior and middle temporal regions perform bottom-up processing while inferior frontal areas send top-down feedback to temporal areas to support lexical-semantic processing (Engel et al., 2001; Badre et al., 2005; Badre and Wagner, 2007; Lau et al., 2008). In neurophysiological studies, these kinds of directional or “causal” influences are often characterized as effective connectivity. However, due to methodological challenges of causality estimation from neuroimaging data, testing hypotheses regarding the interregional influences during N400 generation has remained difficult.

A small number of previous N400 studies have estimated fronto-temporal directional influences using a model-driven method known as Granger causality (Cope et al., 2017; Schoffelen et al., 2017). The Granger causality analysis tests whether information from the past activity of one region can predict future activity in another better than its own past using single variable auto- regressive models (Granger and Hatanaka, 1964). For example, in an MEG experiment with word reading task, the Granger causality method identified inferior frontal cortex and anterior temporal regions to receive widespread input and middle temporal regions to send widespread output (Schoffelen et al., 2017) . In parallel, bi-directional Granger-causal relationships were observed between temporal and frontal sources in matching between degraded spoken words with the previously shown visual word (Cope et al., 2017). However, the limitation of model-driven approaches such as Granger causality, or its analogue “dynamic causal modeling”, is that they require assumptions of the temporal and spatial covariance of the sources, which are difficult to estimate in the presence of noise and with a limited amount of data.

Here, to address the critical barriers on causality modeling, we therefore implemented a novel data-driven approach to estimate the causal connections between frontal and temporal areas during N400 generation. We used data from a classic N400 paradigm with simple German sentences where the final noun was highly predicted (HP) or less predicted (LP) by the previous verb. Our method is based on the temporal generalization technique (King and Dehaene, 2014), which uses machine learning. In this method a classifier is trained on one cortical area’s activity and each time point to discriminate between HP and LP. This classifier is then tested in another cortical area across all time points following temporal generalization idea.

This method allows us to quantify that how much information from one area is predictive of activity in another area and future time points. Our method’s concept is, thus, similar to Granger causality in principle but it is fully data driven and based on multivariate analysis. We tested this method in the context of a study pursuing better understanding of the complex dynamics of feedback and feedforward information flow within fronto-temporal language network during auditory perception of speech.

## Materials and Methods

### Participants

In total, twenty-one right-handed German native speakers (11 female) participated in a MEG experiment (age range: 20-32 years, median: 27) (Oldfield, 1971). The participants reported having no hearing deficits or neurological diseases and they gave a written informed consent before the experiment. The study was conducted in accordance with the declaration of Helsinki, and it received ethical approval from the ethics commission of the University of Leipzig (Ref. 059-11-07032011). Other, more rudimentary aspects of this data set have been analyzed and published in (Maess et al., 2016; Mamashli et al., 2019a).

### Stimuli

Stimuli were short German sentences *[e*.g. *He drives the <u>car</u>, German: Er fährt das <u>Auto</u>*], which were grouped based on the cloze probability values of the nouns. Cloze probability is the probability that mother tongue speakers would select this word to complete the given context (Taylor, 1953). Nouns with a cloze probability >50% were considered as having high semantic predictability (HP), *[e*.g. *He drives the <u>car</u>, German: Er fährt das <u>Auto]</u>* and those with a cloze probability <24% were considered to have low semantic predictability (LP), *[e*.g. *He gets the <u>car</u>, German: Er kriegt das <u>Auto</u>*] (Maess et al., 2016).

### Design and procedure

In a dimly lit shielded room, MEG data were measured with a 306 channel Neuromag Vectorview device (Elekta, Helsinki, Finland), at 500 Hz sampling rate using a bandwidth of 160 Hz (Vacuumschmelze Hanau, Germany). Each participant’s individual hearing thresholds were determined for both ears separately using a subpart of one of the sentences. Stimuli were presented at 48 dB sensation level (i.e., above the mean individual hearing threshold). Each experimental session consisted of five recording blocks. All stimuli were randomized and presented in the first two blocks. Using a different randomization, stimuli were repeated in blocks three and four. The onsets of all sentences, and the onsets of the verbs and the nouns were specifically marked. To control for the accuracy of the MEG inverse solutions, a sequence of simple tones (200 ms length and 500 Hz pitch) was presented during the fifth block with a stimulus onset asynchrony (SOA) of 2200 ms, at the same loudness level as the sentence stimuli.

During each measurement block, participants were instructed to fixate a visually projected cross, to listen carefully to the presented sentences, and to stay motionless. The fixation cross was presented from 700 ms before onset until 700 ms after the offset of each sentence. To keep participants engaged with the listening, in 15% of the sentences, the same or an alternative sentence, was spoken by a male voice. Participants’ (incidental) were asked to judge whether the two preceding sentences (female and male voice) were the same by a button press. A symbolic face was provided to inform participant’s response-to-button-alignment: one happy and one sad face, presented on the left and right side of the screen. Participants answered “yes” with pressing the button at the side of the happy face and “no” with the other using their thumb. The symbolic faces was randomly presented on right or left and counterbalanced over all stimuli in each block.

### Data preprocessing

Signal space separation (SSS) method was used to suppress environmental interference of the MEG data (Elekta-Neuromag Maxfilter software) (Taulu et al., 2004; Taulu and Simola, 2006) and also to transform the data from each block into the same head position (Taulu et al., 2004). To suppress cardiac and eye artifacts, signal space projection was used (Gramfort et al., 2014). Data were extracted into single trials lasting 1.4 seconds, ranging from 400 ms before noun onset to 1000 ms following it. MEG data were filtered with a low pass filter of 25 Hz using MNE-C (fft-based filter) and a highpass of 0.5 Hz with a filter size of 8192. Epochs were rejected if the peak-to-peak amplitude exceeded 150 μV, 1 pT/cm, and 3 pT in any of the electrooculogram, gradiometer, and magnetometer channels, respectively. To equalize the signal-to-noise ratio in each condition (i.e., HP and LP), the number of trials in the lesser populated condition was used to analyze both conditions. The median of the used trials was 97.5 and the minimum number of trials was 79.

### Source estimation

Each participant’s cortical surface representation was reconstructed from 3D structural MRI data using FreeSurfer (http://surfer.nmr.mgh.harvard.edu). The cortical surface was decimated to a grid of 10242 dipoles per hemisphere, i.e., with approximate spacing of 5 mm between adjacent source locations on the cortical surface. The MEG forward solution was computed using a single-compartment boundary-element model (BEM) assuming the shape of the intracranial space (Hämäläinen and Sarvas, 1987). The inner skull surface triangulations was generated from the T1- weighted MR images of each participant with the Freesurfer “wastershed” algorithm. The cortical current distribution was estimated using a depth-weighted, minimum-norm estimate (MNE) (http://www.martinos.org/martinos/userInfo/data/sofMNE.php (Lin et al., 2006)) assuming a fixed orientation of the source, perpendicular to the individual cortical mesh. The noise-covariance matrix used to calculate the inverse operator was estimated from data collected from empty room recordings prior and following the recordings with each subject. To reduce the bias of the MNEs towards superficial currents, we used depth weighting. In other words, the source covariance matrix was adjusted to favor deep source locations.

### Inter-subject cortical surface registration for group analysis

Each participant’s inflated cortical surface was registered to an average cortical representation (FsAverage in FreeSurfer) by optimally aligning individual sulcal-gyral patterns computed in FreeSurfer (Fischl et al., 1999a). To provide more accurate inter-subject alignment of cortical regions than volume-based approaches, we used a surface-based registration technique based on folding patterns (Fischl et al., 1999b; Van Essen and Dierker, 2007).

### Region identification and analysis

The analysis were focused on six cortical areas of the FreeSurfer Desikan-Killiany Atlas in both hemispheres, which are believed to constitute the most critical parts of semantic language networks (Lau et al., 2008; Price, 2010; Friederici, 2011), including bilateral superior temporal gyrus (STG), middle temporal gyrus (MTG), and inferior frontal gyrus (IFG) including Brodmann areas BA44, BA45 and BA47. In addition, we used an automatic routine (mris_divide_parcellation) available in the Freesurfer package (equal size principle) to break each large region into smaller equal size sub-regions; i.e., all sub-regions in all regions were of approximately the same size—thereby increasing the spatial specificity for further analysis (Mamashli et al., 2017; Mamashli et al., 2019b; Mamashli et al., 2019c; Mamashli et al., 2020; Mamashli et al., 2021a; Mamashli et al., 2021b), as areas can lead to temporal signal cancellations. Furthermore, we grouped the sub-regions into anterior and posterior parts of each cortical region, e.g., STG will be divided into anterior STG (aSTG) and posterior STG (pSTG). In total, we had six regions of interest (ROI) in each hemisphere: aSTG, pSTG, aMTG, pMTG, aIFG and pIFG (**Figure 1**).

**Figure 1:**
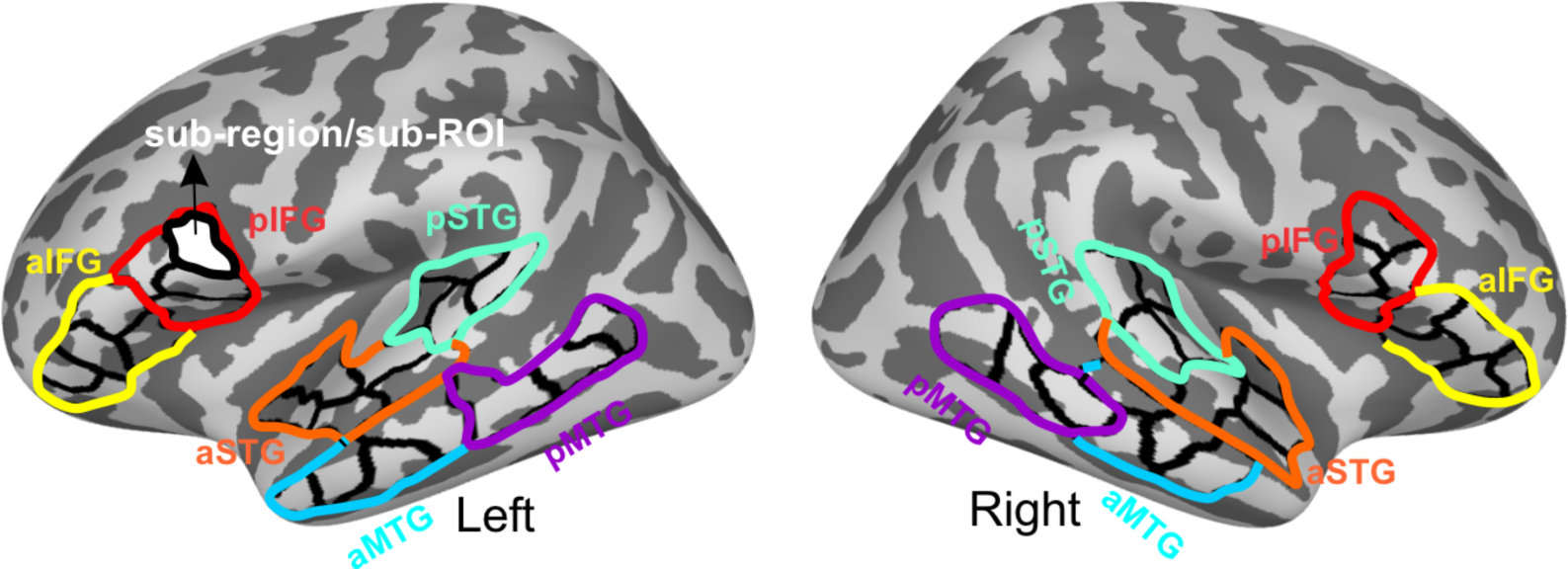
ROIs and sub-regions or sub-ROIs in left and right hemisphere.

### Sub-region time series extraction

Epochs were extracted and averaged across all vertices within each sub-region, to compute the mean sub-region time course, generating ***X***(Λ, *T*, *N*), where Λ is the number of vertices, T is the number of time points, and *N* is the number of epochs. Since the individual vertex (dipole) orientations is ambiguous, these time series were first aligned with the dominant component of the multivariate source time course, and then averaged to calculate the sub-region mean. In order to align the sign of the time series across vertices, we first concatenate all the epochs for each vertex in a single time series and then computed an SVD of the data **X**^T^ = **UΣV**^**T**^. The sign of the dot product between the first left singular vector U and all other time-series in a sub-region was computed. If this sign was negative, we inverted the time-series before averaging over all time courses of a sub-region. Finally, temporal data of each sub-region was arranged as a 2D matrix [epochs X time].

### Inter-regional temporal generalization Multivariate Pattern Analysis (MVPA)

Here, we use a data-driven multivariate approach to estimate the causal connection between two regions. Multivariate pattern analysis has been used before both using MEG (King and Dehaene, 2014; Cichy and Pantazis, 2017; Mohsenzadeh et al., 2018; Hatamimajoumerd et al., 2020) and fMRI (Hatamimajoumerd et al., 2022), where a classifier is trained in one experimental condition and tested in another condition. In contrast, here, a classifier is trained to learn the difference between conditions at one point of time in one region and then tested in at another point of time in another region.

To accomplish this, an SVM classifier is trained across two conditions (LP vs HP) in ROI_1_ using the sub-ROI activities as features at each time point. This classifier is then tested in ROI_2_ and across all time points using temporal generalization idea. This process is replicated for all time points of ROI_1_ and eventually provides the temporal generalization matrix for each ROI pair. The time window was from -200ms to 800ms. We focused our analysis on the within hemisphere ventral and dorsal path in language processing to investigate the information flow from anterior IFG to anterior temporal areas (e.g., aIFG to aSTG/aMTG) and posterior IFG to posterior temporal (e.g., pIFG to pSTG/pMTG).

Similarly, we tested the opposite direction from temporal to IFG (e.g., pSTG/pMTG to pIFG). For simplicity, we refer these patterns to as “directional connections”. In total, we tested eight directional connections in each hemisphere. To increase the signal-to-noise ratio, we randomly selected 10 epochs, averaged within each condition, and bootstrapped this 100 times (Cichy and Pantazis, 2017). A schematic display of the method is shown in **Figure 2**.

**Figure 2:**
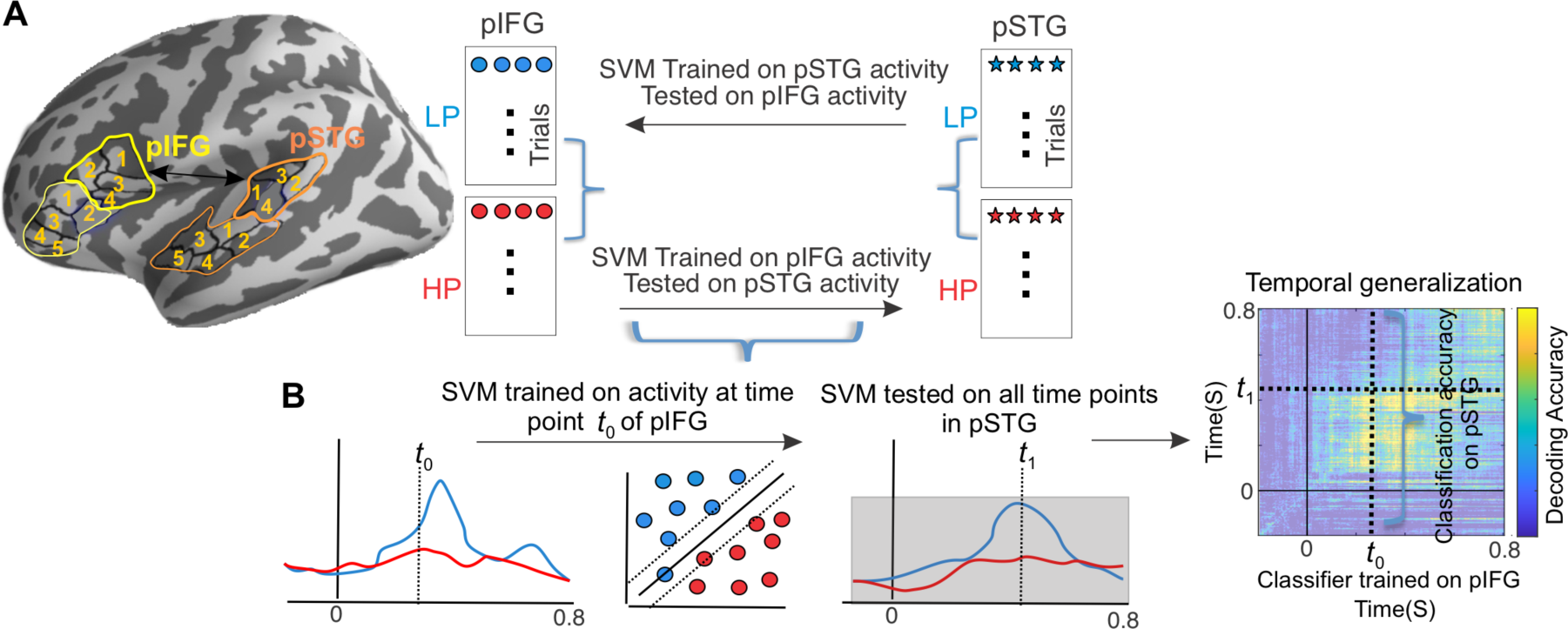
A schematic display of the method. (**A**) Examples of ROIs and sub-ROIs in pIFG and pSTG. SVM classifier is trained on four features from four sub-ROIs activity in pIFG to classify LP from HP conditions and then tested on the four features extracted from pSTG sub-ROI activity. Similarly, the same process was done from pSTG to pIFG. (**B**) SVM classifier is trained at each time point of pIFG activity and tested on all time points of pSTG. The accuracy of model from pSTG test data is used to create temporal generalization matrix. Here, one time point t_0_ and t_1_ are shown as an example.

### Statistical analysis

For each directional connectivity between a pair of ROIs, cluster-based statistics were applied (Maris and Oostenveld, 2007). We used P < 0.025 as the initial threshold, 1000 permutations, and one-tailed one-sample t-tests as the test statistics against the chance level for binary classifier. We estimated the empirical chance level using simulations by shuffling the labels 100 times and performed the temporal generalization for all subjects and connections. The temporal generalization matrix was flattened and gained 10000 shuffled accuracies. To generate a null distribution, values were pooled across all subjects and connections. The null distribution was Gaussian with mean at 0.5. Therefore, the empirical chance level for our case was 0.5. Thus, we used 0.5 as the chance level in our test statistic. In addition, to correct for 16 directional connectivity tests, we applied false discovery rate (FDR) method at 0.025 thresholds. The 0.025 thresholds were chosen to account for the one-tailed t-test. For each connection, we considered the first 3 clusters as they represent the strongest effect. In summary, we applied FDR on 16 × 3= 48 tests.

## Results

### Information flow from temporal areas to IFG

We tested for across-areal generalization by training the classifier on MTG and STG evoked response activity and then testing this classifier on IFG activity. Any cluster above the diagonal shows how earlier time in temporal areas affect future time in frontal, which we interpret as reflecting feedforward-type influences. We observed this pattern in five out of 6 significant connections (**Figure 3**). These patterns included from influences from the left pSTG to left pIFG, left pMTG to left pIFG, left aMTG to left aIFG, right pSTG to right pIFG, and right pMTG to right pIFG. The left pSTG influenced pIFG processing at multiple time intervals started from 250ms and extended later to 500ms (**Figure 3A**). From the left pMTG to pIFG and from the left aMTG to aIFG, there was a continous feedforward effect from 50ms to 450 and 500ms respectively (**Figure 3C-D**). In the case of connectivity patterns from the right pSTG to pIFG and from the right pMTG to pIFG, the feedforward effects were more discontinous (**Figure 3E-F**).

**Figure 3:**
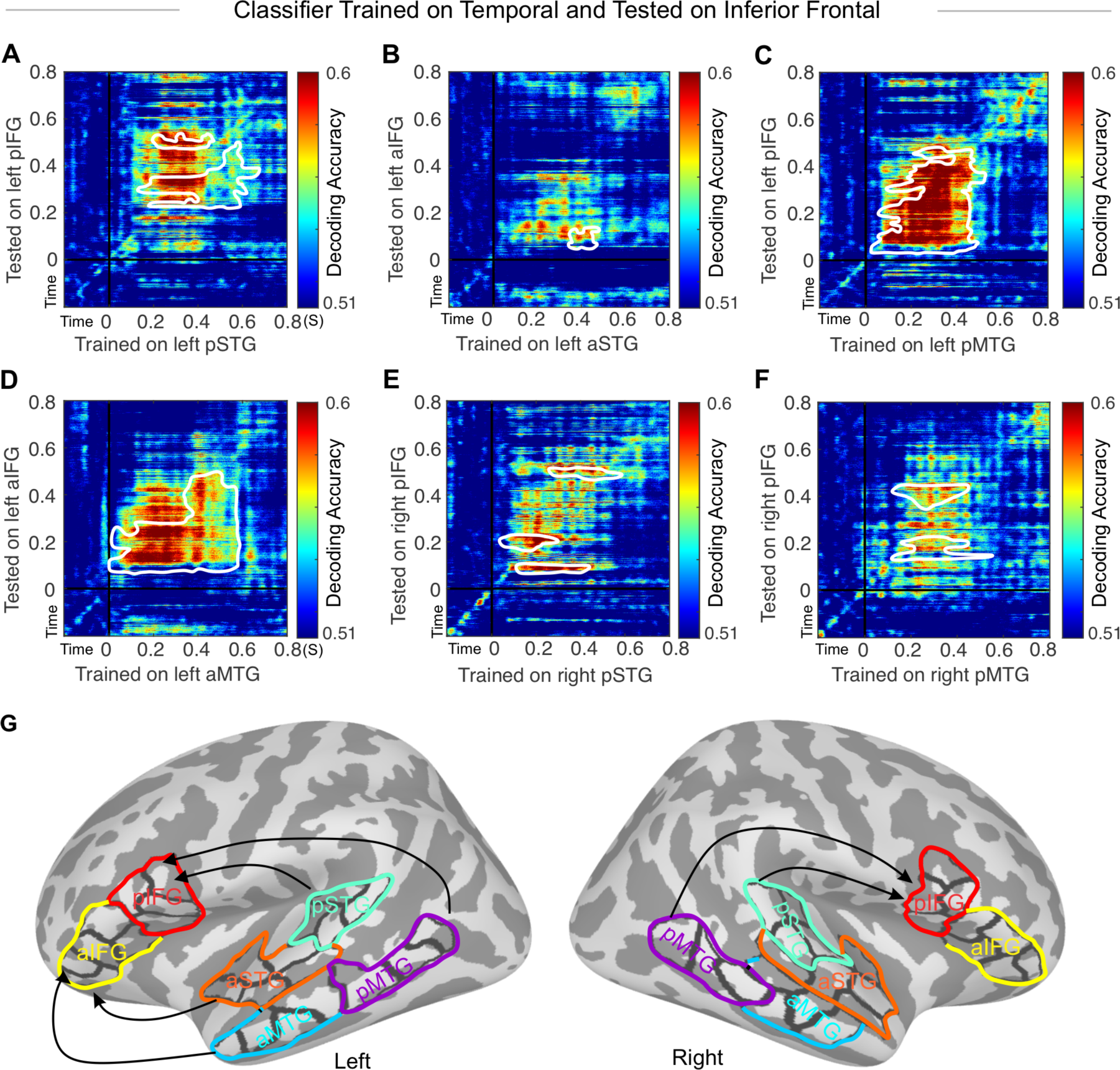
Temporal generalization decoding matrix averaged over all subjects. The white contour indicates significant decoding values against the chance level. SVM classifier is trained on (A) left pSTG and tested on left pIFG, (B) left pMTG and tested on left pIFG, (C) right aSTG and right aIFG, and (D) right pMTG and right pIFG.

### Information flow from IFG to temporal areas

Analogously to the above section, we tested for across-areal generalization by training the classifier on IFG evoked response activity and then testing this classifier on STG and MTG activity. We found significantly larger than chance level (50%) accuracy in 6 connections corrected for multiple comparisons (**Figure 4**). These included left aIFG to left aSTG, left aIFG to left aMTG, left pIFG to left pMTG, right pIFG to right pMTG, right pIFG to right pSTG, and right aIFG to right aSTG. The temporal generalization dynamics were different in each connection. When the cluster expands above the diagonal, it shows that at each time, the classifier trained in IFG is predictive of future time points in temporal-cortex areas. We interpret these kinds of patterns as reflecting feedback influences from IFG to temporal areas. From left aIFG to left aSTG and aMTG, there were effects up to 250ms and 450ms respectively that started as early as 50ms and were sustained for at least 200ms (**Figure 4A-B**). The effect from left pIFG to pMTG started later around 250ms and continued for about 200ms and affected time interval after 450ms (**Figure 4C**). The earliest frontal effect seemed to start from right pIFG to pMTG and pSTG (**Figure 4D-E**), where the influence on pMTG lasted longer time up to 400ms, whereas in pSTG up to around 150ms. There was also a continuous effect from right aIFG to aSTG around 200ms for a short duration. Furthermore, from right aIFG to aSTG, there was a small effect before 200ms.

**Figure 4:**
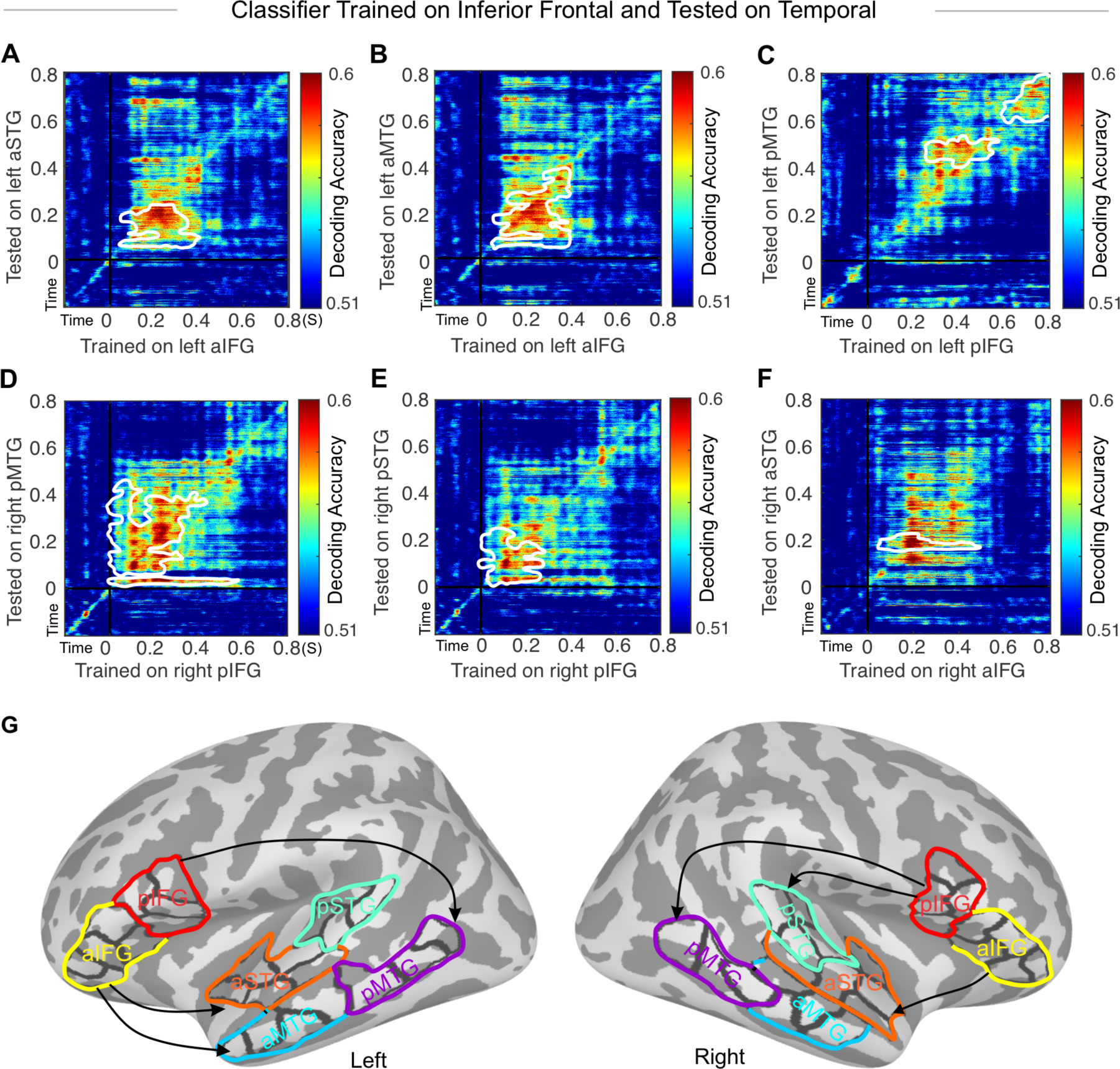
Temporal generalization decoding matrix averaged over all subjects. The white contour indicates significant decoding values against the chance level. SVM classifier is trained on (A) left pIFG and tested on left pSTG, (B) left pIFG and tested on left pMTG, (C) left aIFG and tested on left aSTG, (D) left aIFG and tested on left aMTG, (E) right aIFG and tested on right aSTG, and (F) right aIFG and tested on right aMTG. (G) The ROIs and the significant connections from A to F are displayed in a cortical surface representation.

Those connections that are significant in both directions are plotted side-by-side in Figure S1 for a better comparison.

## Discussion

In this study, we used a novel approach to investigate feedforward and feedback influences between inferior frontal and temporal cortex areas, using source estimates of event-related MEG responses to low-predictability vs. high-predictability nouns.

Our rationale rests on the utility of temporal generalization methods in multivariate classification of brain data from specific brain areas: The idea is that when the classifier performance exceeds chance level in the testing area at future time points, this means that brain activity in the training area contain information that helps predicting the LP vs HP condition future time points in the testing area. By examining instances when the classifier training was based at an earlier time period than the testing, we made inferences on potential directional influences in language processing underlying N400 generation.

We here have presented evidence for both feedforward influences from STG/MTG to IFG and feedback influences from IFG to STG/MTG in both hemispheres. The strongest feedforward effects were observed from the left pSTG/pMTG to the left pIFG and from the left aMTG to the left aIFG. In parallel, strongest feedback effects were from the left aIFG to the left aSTG/aMTG and from the right pIFG to the right pSTG/pMTG. These results provide evidence that feedforward and feedback influences are transferred through both ventral and dorsal pathways and they are not restricted to a certain path. Dorsal and ventral are the two main structural pathways for language processing. The ventral pathway connects the temporal cortex to inferior frontal regions via extreme fiber capsule system (EFCS) and uncinate fascile (UF) and the dorsal pathway connects the posterior frontal area to posterior part of the temporal cortex via arcuate fascile (AF) and the superior longitudinal fascicle (SLF) (Friederici, 2012). Moreover, our results suggest that feedforward influences are mostly left lateralized whereas feedback influences are present in both hemispheres. In our previous study (Maess et al, 2016), focused on the evoked responses of the verbs and the nouns, we observed a reduction of the N400 response for highly predicted nouns as expected and the opposite pattern for the noun-preceding verbs. Highly predictive verbs yielded stronger N400 amplitude compared to less predictive verbs. Enhanced activity for highly predictive relative to less predictive verbs, were interpreted to reflect pre-activations of semantic features associated with the expected nouns. Therefore, it is interesting that feedback influences start at a very early latencies, almost immediately after the stimulus onset. In contrast, the majority of feedforward influences started at least with a 100 ms delay.

A number of competing models have been proposed on top-down and bottom-up influences between temporal and frontal areas during sentence comprehension (Friederici 2012). Verifying such models has been difficult due to the complications in estimating causality in human recordings. Indeed, to date, only a few previous studies have estimated the causal connections between temporal and frontal areas in predictive speech processing using more classic techniques. Cope et al. (2017) found bi-directional fronto-temporal causal connections using Granger causality in distinct frequency bands when spoken words were matched with visual presentation. Using similar method in a reading task, Schoffelen et al. (2017) found feedforward connection from pMTG to IFG and feedback and feedforward connections between IFG and aMTG. A recent study (Schroen et al., 2023) using a subset of our stimulus material investigated temporo-frontal causal influences with a combined transcranial magnetic stimulation and electroencephalography approach. Interestingly, using this completely different approach, they also observed early feedforward influences from left pSTG to left IFG and late feedback influences (300-500ms) from left IFG to left pSTG. Consistent with these previous results, the present results highlight the importance of bidirectional interactions between functionally specialized brain regions to facilitate complex language processing (Friederici 2012). Our novel inter-regional temporal generalization could facilitate quantitative testing of theoretical models proposed for language processing in general (Hickok and Poeppel, 2004, 2007; Friederici, 2011) and N400 processing in particular (Lau et al., 2008).

Estimating feedback and feedforward influences using neuroimaging data has been challenging. One of the inherent properties of these connections is that feedforward influence is time-locked and stimulus-driven, whereas feedback influences associated with cognitive processing can be presumed to jitter in time and vary more prominently subject by subject, for example, due to individual differences in cognitive capacities. The more pronounced variability within and between individuals weakens the estimated representations of feedback influences in time relative to feedforward influences, making their quantitative estimation harder. This could be one of the factors why in the present study, the average decoding accuracy was stronger in feedforward than feedback connections.

## Conclusion

In summary, we implemented a novel method to estimate feedback and feedforward influences using cross-regional temporal generalization in MEG decoding. Aiming to understand the information flow in N400 generation in a simple language paradigm, we found IFG feeding back to STG/MTG bilaterally and STG/MTG feeding forward to IFG left-lateralized. Our results are consistent with the long-standing but empirically challenging notion that dynamic feedback and feedforward influences between IFG and temporal areas drive N400 generation.

## Acknowledgements

This research was supported by the Max Planck Society, the International Max Planck Research School Leipzig and by NIH grants: R01DC016915, R01DC016765, R01DC017991, P41EB015896. We thank Yvonne Wolff for help in acquiring the data.

## Data and Code Availability

Data and code will be available upon request.

## Author Contributions

F.M. conceived the study, designed the study, conducted the experiments, analyzed the data, and wrote the manuscript. S.K., J.A., conceived the study, designed the study, analyzed the data, and wrote the manuscript. A.F., B.M., and J.O. designed the study, conducted the experiments, and wrote the manuscript, E.H., M.J., I.U, K.L, analyzed the data and wrote the manuscript.

## Supplementary materials

**Figure S1:**
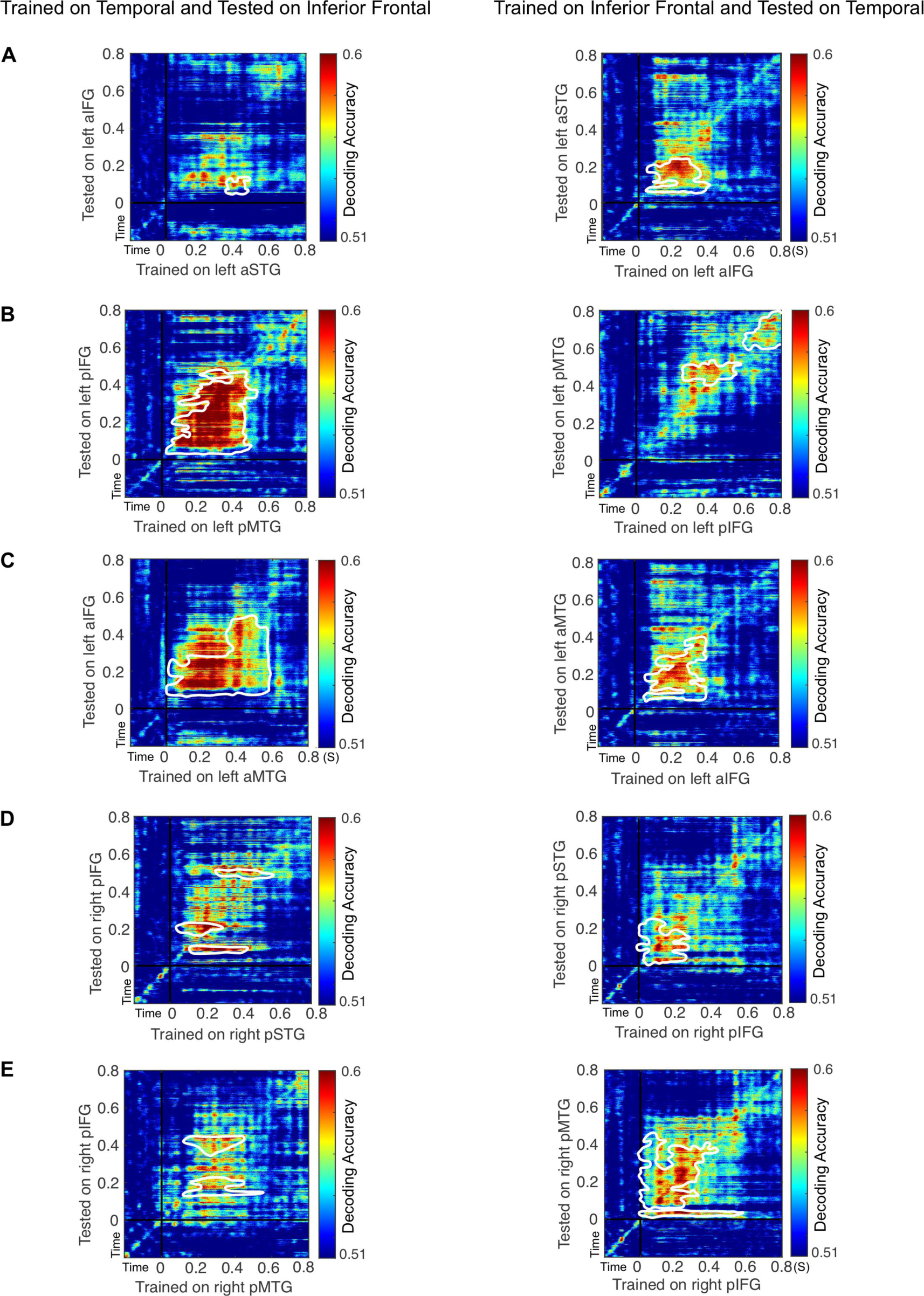
Significant connections that are bi-directional. The left panel shows the feedforward effects and the right panel is the feedback effects.

